# The sequence of regional structural disconnectivity due to multiple sclerosis lesions

**DOI:** 10.1101/2023.01.26.525537

**Authors:** Ceren Tozlu, Emily Olafson, Keith Jamison, Emily Demmon, Ulrike Kaunzner, Melanie Marcille, Nicole Zinger, Nara Michaelson, Neha Safi, Thanh Nguyen, Susan Gauthier, Amy Kuceyeski

**Affiliations:** Department of Radiology, Weill Cornell Medicine, New York, NY, USA; Department of Neurology, Weill Cornell Medical College, New York, New York, USA

**Keywords:** multiple sclerosis, disease progression, cognition, chronic active lesions, event-based modeling

## Abstract

**Objective:** Prediction of disease progression is challenging in multiple sclerosis (MS) as the sequence of lesion development and retention of inflammation within a subset of chronic lesions is heterogeneous among patients. We investigated the sequence of lesion-related regional structural disconnectivity across the spectrum of disability and cognitive impairment in MS.

**Methods:** In a full cohort of 482 patients, the Expanded Disability Status Scale was used to classify patients into (i) no or mild vs (ii) moderate or severe disability groups. In 363 out of 482 patients, Quantitative Susceptibility Mapping was used to identify paramagnetic rim lesions (PRL), which are maintained by a rim of iron-laden innate immune cells. In 171 out of 482 patients, Brief International Cognitive Assessment was used to identify subjects with cognitive impairment. Network Modification Tool was used to estimate the regional structural disconnectivity due to MS lesions. Discriminative event-based modeling was applied to investigate the sequence of regional structural disconnectivity due to all representative lesions across the spectrum of disability and cognitive impairment.

**Results:** Structural disconnection in the ventral attention and subcortical networks was an early biomarker of moderate or severe disability. The earliest biomarkers of disability progression were structural disconnections due to PRL in the motor-related regions. Subcortical structural disconnection was an early biomarker of cognitive impairment.

**Interpretation:** MS lesion-related structural disconnections in the subcortex is an early biomarker for both disability and cognitive impairment in MS. PRL-related structural disconnection in the motor cortex may identify the patients at risk for moderate or severe disability in MS.

## INTRODUCTION

The size, location, and underlying pathology of white matter (WM) lesions are heterogeneous among people with multiple sclerosis (MS)^1^, making the prediction of disease progression challenging. Moreover, the sequence in which lesions disrupt various brain regions’ structural connectivity (WM streamlines) and how these disruptions impact disability progression and/or cognitive impairment in MS has not yet been studied. Discriminative event-based modeling (dEBM) is an approach used to estimate the temporal sequence of biomarker abnormalities over disease progression using cross-sectional data^2–7^. This data-driven approach has been used to investigate the sequence of gray matter (GM) atrophy only^2^, or the sequence of changes in a combination of features like atrophy, lesion load, properties of the brain’s functional networks, and microstructure of major white matter tracts^3^ across the stages of motor and cognitive impairment in MS. However, these studies restricted the structural biomarkers by using a-priori defined regions of interest; here we begin with whole-brain biomarkers selected in a data-driven manner.

Motor and cognitive impairments in MS have been related to disruptions in the network of WM tracts in the brain, referred to collectively as the brain’s structural disconnectivity^4,5^. The Network Modification (NeMo) Tool^6^, which measures structural disconnectivity due to lesions, has been used by our group and others to correlate lesion-related structural disconnectivity patterns to impairments, outcomes, functional connectivity disruptions, rehabilitation response, and GM pathology in people with MS (pwMS)^4,7–10^. Importantly, our recent study of pwMS demonstrated the clinical utility of the NeMo Tool in understanding mechanisms of disease by showing that the structural disconnectivity estimated using the NeMo Tool was more predictive of disability than structural connectivity measured using an individual’s diffusion MRI (dMRI)^8^.

In addition to their size and location, a lesion’s underlying pathology can also influence its impact on disability. Chronic CNS inflammation in the MS lesion is maintained, in part, with iron-laden pro-inflammatory microglia/macrophages at the rim of chronic active MS lesions^11–13^. Quantitative Susceptibility Mapping (QSM)^14,15^ has been used to identify paramagnetic rim lesions (PRL)^16^ and has expanded our understanding of chronic active lesions on disease mechanisms and disease progression^16–19^. Structural disconnectivity due to PRL has been shown to be better associated with disability as compared to structural disconnectivity due to non-PRL^5^. However, it remains unknown if the sequence of regional structural disconnectivity differs among PRL compared to non-PRL and if this difference impacts clinical disability.

Here we use dEBM to investigate the sequence of regional structural disconnectivity due to all MS lesions (as identified on T2 FLAIR), PRL, and non-PRL, estimated using the NeMo Tool, as disability and cognitive impairment range from less to more severe. Understanding the impact of the lesion size, location, and classification on multiple clinical outcomes may provide an opportunity to identify the patients at risk for future disability and allow personalized treatments to minimize the burden of MS.

## MATERIAL AND METHODS

### Subjects

Four hundred-eighty-two pwMS (age: 41.83 ± 11.63 years, 71.57% females, the “ ± ” sign indicates standard deviation) with a diagnosis of clinically isolated syndrome (CIS) (n=16) or MS (430 relapsing-remitting and 36 primary or secondary progressive MS) were enrolled in our study; inclusion criteria included no contraindications to MRI. Demographic data was collected (age, sex, clinical phenotype, and disease duration) and subjects underwent MRI. Extended Disability Status Score (EDSS) was used to quantify disability, where an EDSS < 3 was considered no and mild disability and EDSS ≥3 was considered moderate to severe disability ^20^. All studies were approved by an ethical standards committee on human experimentation and written informed consent was obtained from all patients.

The Brief International Cognitive Assessment for Multiple Sclerosis (BICAMS) was used to assess cognition in a subset of 171 pwMS. BICAMS consists of three assessments: 1) The Symbol Digit Modalities Test (SDMT), which measures the processing speed, 2) California Verbal Learning Test-II (CVLT-II) Immediate Recall (Total of Trials 1-5), which measures verbal learning and short-term memory, and 3) Brief Visuospatial Memory Test-Revised (BVMT-R) Immediate Recall (Total of Trials 1-3), which measures visuospatial learning and short-term memory^21^. A higher score of each test is associated with better cognitive performance. If a subject’s SDMT z-score was <-1 (21% of 171 pwMS) or CVLT or BVMT t-scores were <40 (10% of 171 pwMS for CVLT and 23% of 171 pwMS for BVMT), they were considered significantly impaired on that test^21^. In this paper, we identified pwMS to be cognitively impaired (CI) if they showed impairments in at least two cognitive tests; otherwise, they were considered cognitively preserved (CP)^22^.

### Image acquisition, processing, and connectome extraction

MRI images were obtained on two different scanners:

1. *<u>The Siemens scanning protocol consisted of the following sequences</u>*: 1) 3D sagittal T1-weighted (T1w) MPRAGE: Repetition Time (TR)/Echo Time (TE)/Inversion Time (TI) = 2300/2.3/900 ms, flip angle (FA) = 8°, GRAPPA parallel imaging factor (R) = 2, voxel size = 1.0 × 1.0 × 1.0 mm3; 2) 2D axial T2-weighted (T2w) turbo spin echo: TR/TE = 5840/93 ms, FA = 90°, turbo factor = 18, R = 2, number of signal averages (NSA) = 2, voxel size = 0.5 × 0.5 × 3 mm3; 3) 3D sagittal fat-saturated T2w fluid attenuated inversion recovery (FLAIR) SPACE: TR/TE/TI = 8500/391/2500 ms, FA = 90°, turbo factor = 278, R = 4, voxel size = 1.0 × 1.0 × 1.0 mm3.
2. *<u>The GE scanning protocol consisted of the following sequences:</u>* 1) 3D sagittal T1w BRAVO: TR/TE/TI = 8.8/3.4/450 ms, FA = 15°, voxel size = 1.2 × 1.2 × 1.2 mm3, ASSET parallel imaging acceleration factor (R) = 1.5; 2) 2D axial T2w fast spin echo: TR/TE = 5267/86 ms, FA = 90°, echo train length = 100, number of excitations (NEX) = 2, voxel size = 0.6 × 0.9 × 3.0 mm3; 3) 3D sagittal T2w FLAIR CUBE: TR/TE/TI = 5000/139/1577 ms, FA = 90°, ETL = 162, R = 1.6, voxel size = 1.2 × 1.2 × 1.2 mm3.

Both scanners had similar parameters for the *<u>axial 3D multi-echo GRE sequence for QSM</u>*: axial field of view (FOV) = 24 cm, TR/TE1/ΔTE = 48.0/6.3/4.1 ms, number of TEs = 10, FA = 15°, R = 2, voxel size = 0.75 × 0.93 × 3 mm3, scan time = 4.2 min using a fully automated Morphology Enabled Dipole Inversion (MEDI+0) method zero-referenced to the ventricular cerebrospinal fluid^23^. The QSM protocol has been harmonized for both scanner manufacturers and was demonstrated to be reproducible across manufacturers^24^.

### Lesion mask creation

The conventional images (T1w, T2w, T2w FLAIR) were co-registered to the sum-of-squares echo-combined magnitude GRE images using the FMRIB’s Linear Image Registration Tool algorithm ^25^; automated brain segmentation was performed using FreeSurfer ^26^. WM tissue segmentations were manually edited for misclassification due to WM T1-hypointensities associated with lesions. The WM hyperintensity lesion masks were created from the T2 FLAIR images by categorizing the tissue type based on the image intensities within the Lesion Segmentation Tool (LST) using the LPA method; the masks generated were further hand-edited if necessary. Next, the T2FLAIR lesions masks were coregistered to the QSM images and further hand edited (if needed) to better match the lesion geometry on QSM. A lesion was designated as PRL if QSM was hyperintense at the edge of the lesion in either a complete or partial manner. The presence of a partial and/or full hyperintense QSM rim (i.e. PRL) was determined as a consensus of two trained reviewers and in the case of disagreement, an independent third reviewer decided on the presence of a positive hyperintense rim. Once the PRL were identified, they were removed from the T2FLAIR lesion masks to obtain a non-PRL mask.

Lesion masks were transformed to the individual’s T1 native space using the inverse of the T1 to GRE transform and nearest neighbor interpolation. Individual T1 images were then normalized to 1mm MNI152 v6 space using FSL’s linear (FLIRT) and non-linear (FNIRT) transformation tools (http://www.fmrib.ox.ac.uk/fsl/index.html); transformations with nearest neighbor interpolation were then applied to transform the native anatomical space T2FLAIR lesion masks to MNI space. The transformations were concatenated (T2FLAIR to T1 to MNI) to minimize interpolation effects. Lesions were visually inspected after the transformation to MNI space to verify the accuracy of coregistration. QSM was available in 363 pwMS to allow for designation of individual chronic lesions as a PRL or non-PRL. Forty three percent (156/363) of these patients had ≥ 1 PRL, therefore separate PRL and non-PRL (T2 FLAIR lesions minus PRL) lesion masks were created for these subjects.

### Network Modification Tool

The MNI space T2 FLAIR lesions (for all subjects), PRL (for the subjects who had ≥ 1 PRL) and non-PRL (for the subjects who had ≥ 1 PRL) masks were processed through the newest version of the Network Modification (NeMo) Tool ^27^, NeMo Tool 2.0, that estimates a lesion mask’s subsequent pattern of structural disconnectivity. The pattern of structural disconnectivity metric was defined for 68 cortical (from the Desikan-Killany atlas), and 18 subcortical and cerebellar regions as the percent of tractography streamlines connecting that region that also pass through the lesion mask. The newest version of the tractography database consists of structural connectomes from 420 unrelated healthy controls (206 female, 214 male, 28.7 ± 3.7 years)^8^. Structural disconnectivity metrics were calculated separately using PRL and non-PRL masks as well as for the complete T2 FLAIR lesion masks. The structural disconnectivity metrics from the T2 FLAIR lesion mask were computed across all subjects (N=482), while the structural disconnectivity metrics scores from PRL vs non-PRL masks were computed only for the subjects that had QSM imaging (N=363). Those subjects having QSM imaging but no PRL were assigned PRL structural disconnectivity scores of 0, since there was no structural disconnectivity due to PRL for these subjects.

### Discriminative event-based modeling

We used discriminative event-based modeling (dEBM) applied to cross-sectional data to estimate the most likely sequence in which regions appear increasingly disconnected across the spectrum from no/mild disability to moderate/severe disability or from cognitively preserved (CP) to cognitively impaired (CI) ^28,29^ (see Figure 1). The conversion of a metric (in this case regional structural disconnection) from healthy to pathological is called as event (*E*_*pn*_), where n indicates the subject *n* = 1, 2, …, *N*and *p* indicates the region *p* = 1, 2, …, *P*. First, dEBM estimates the posterior probability of each region being within healthy range (i.e., less disconnected) vs. pathological (i.e., more disconnected) using a Gaussian mixture model (GMM) of the two subject groups’ distributions of regional structural disconnectivities. The GMM allows calculation of the likelihood that region *p* of subject *n* is pathological, i.e. *P*(*x*_*pn*_|*E*_*n*_), or healthy, i.e. *P*(*x*_*pn*_| − *E*_*n*_). Second, the sequence (*S*) of the events (i.e., regions becoming pathologically disconnected) is created by maximizing the following likelihood:

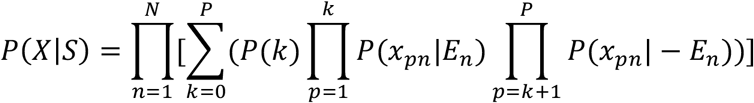

where *X* is the data matrix containing the regional disconnectivity scores for all subjects and *P(k)* is the prior probability of being at stage *k*, which indicates that events *E*_1,_ …, *E*_*k*_ have occurred but *E*_*k*+1,_ …, *E*_*P*_ have not yet occurred. The entire algorithm is repeated using 100 bootstrapped samples of the data to compute the uncertainty (i.e., the positional variance of the sequence) and standard error of the event centers. DEBM results are visualized via a matrix with rows indicating regions and columns indicating position in the sequence of disability/cognitive impairment progression. The intensity of each matrix entry corresponds to the certainty of that region’s structural disconnection at that position in the sequence across 100 bootstraps (darker color with greater certainty). The codes to perform the dEBM analysis are publicly available (https://github.com/EuroPOND/pyebm).

**Figure 1:**
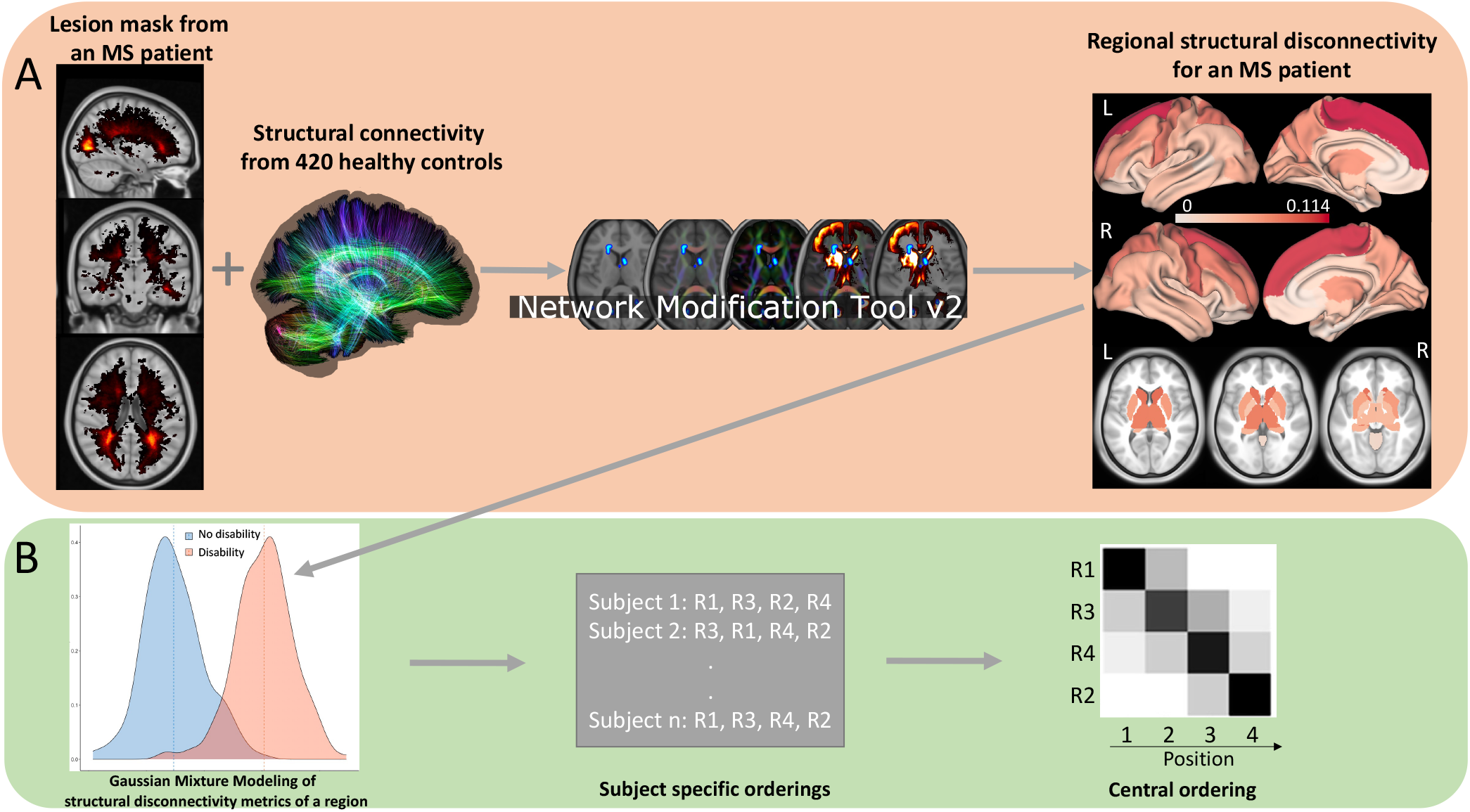
(A) The NeMo tool was used to predict regional structural disconnectivity metrics from the pwMS’s lesion masks. (B) The dEBM algorithm was applied to the regional structural disconnectivity metrics to estimate the sequence of regional structural disconnections as disability/cognitive impairment occurs. First, a Gaussian Mixture Modeling was used to calculate the probability distributions of the regional structural disconnectivity metrics for two groups (for example: no disability vs disability groups). Second, the likelihood defined in the text is maximized by finding the optimal sequence of regional structural disconnection “events”. Results are visualized via a matrix with rows indicating regional structural disconnectivity and columns indicating the position in the sequence of disability/cognitive impairment progression. The intensity of each matrix entry corresponds to the certainty of that region’s structural disconnection at that position in the sequence across 100 bootstraps (darker color means greater certainty in event position).

In our study, we applied the dEBM algorithm to 3 different datasets:

1. Regional structural disconnectivity due to T2 FLAIR lesions in all subjects (N=482) across the spectrum from no or mild disability (EDSS < 3) (N=379) to moderate or severe disability (EDSS ≥ 3) (N=103)
2. Regional structural disconnectivity due to PRL vs non-PRL separately in a subset of the subjects (N=363) across the spectrum from no or mild disability (EDSS < 3) (N=272) to moderate or severe disability (EDSS ≥ 3) (N=91)
3. Regional structural disconnectivity due to T2 FLAIR lesions in a subset of the subjects (N=171) across the spectrum from CP (N=150) to CI (N=21)

For each dataset, we compared the regional structural disconnectivity metrics between disability or cognition groups using the Student’s t-test. Similar to the approach that Eshaghi et al. used^2^, we included the most significantly different 20 regions in the dEBM algorithms to reduce the dimensionality of the model. In addition to the structural disconnectivity metrics, confounding variables such as age and sex were included in all analyses.

### Mass univariate analysis

First, demographics and clinical variables were tested for differences between the no/mild disability vs moderate/severe disability groups and the CP vs CI groups using the Chi-squared test for qualitative variables and Wilcoxon rank-sum test for quantitative variables. Student’s t-tests were used to quantify group differences in the regional structural disconnectivity metrics (after logarithmic transformation to ensure normality). Differences were considered significant when p<0.05 after Benjamini-Hochberg (BH) correction for multiple comparisons^30^. All statistical analyses were performed and graphs were created using R version 3.4.4, Matlab version R2021b, and Python version 3.9.

## RESULTS

### Patient Characteristics

Table 1 shows the demographics, clinical, and imaging features of all subjects and by disability (no/mild vs moderate/severe disability), having PRL (at least one PRL vs no PRL), and cognition (CP vs CI) groups. The pwMS who had moderate to severe disability were older and had greater disease duration compared to those with no to mild disability (p-value <0.05). The pwMS who had at least one PRL were younger and had greater lesion volume compared to the pwMS without PRL (p-value <0.05). Thirty-six, 18, and 40 pwMS had impairments on SDMT, CVLT, and BVMT, respectively. Twenty-one subjects were included in the CI group as these subjects showed impairments in at least two cognitive tests. Age and disease duration did not differ between CP and CI subjects (p-value >0.05), while lesion volume was greater in pwMS with moderate to severe disability compared to those with no to mild disability and in CI compared to the CP group (p-value <0.05).

**Table 1:**
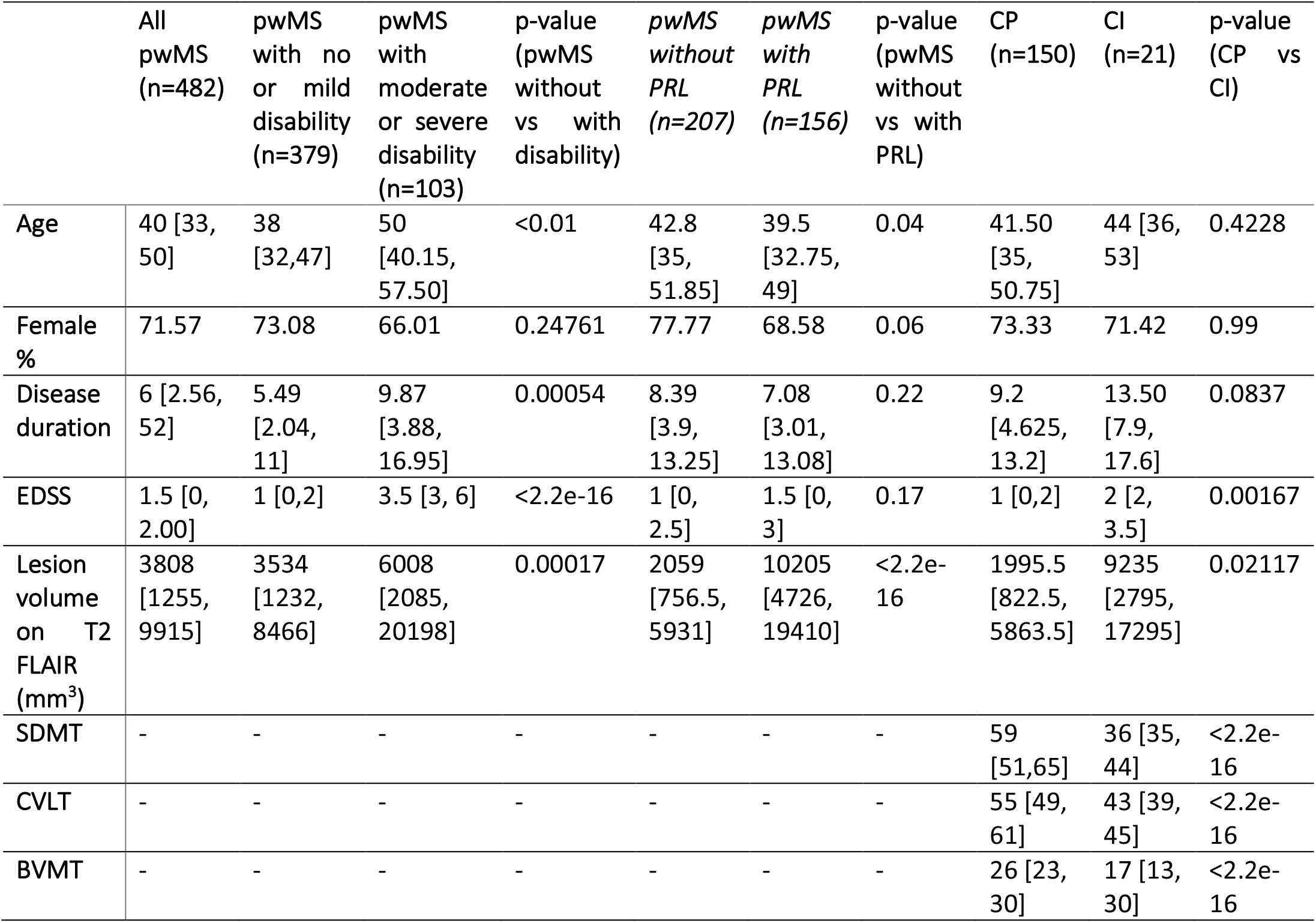
Patient demographics and imaging features for all pwMS and the two sets of subgroups - no/mild vs moderate/severe disability and CP vs CI. The p-values were obtained in the comparison of no/mild vs moderate/severe disability groups (5th column) and CP vs CI (8th column) using the Wilcoxon rank-sum test. Values were presented as median [1st quartile, 3rd quartile] for the continuous variables. P-values were corrected with the BH method for multiple comparisons. Age and disease duration were measured in years.

### Sequence of Structural Disconnection due to MS Lesions Across the Spectrum of Disability

Figure 2 shows the t-statistics of the structural disconnectivity metrics for the 20 regions that were the most significantly different between disability groups and the dEBM model results, visualized as the sequence of the structural disconnectivity metrics across disability severity. All subjects (n=482) were used for this analysis. The structural disconnectivity metrics were significantly greater, particularly in the bilateral precuneus, right postcentral, and right superior parietal regions in the moderate/severe disability group compared to the no/mild disability group (Fig. 2 A). The dEBM results showed that structural disconnectivity due to T2 FLAIR lesions occurs first in the right supramarginal, followed by subcortical regions including putamen, pallidum, cuneus, and thalamus as disability becomes more severe (Fig. 2, B and C).

**Figure 2:**
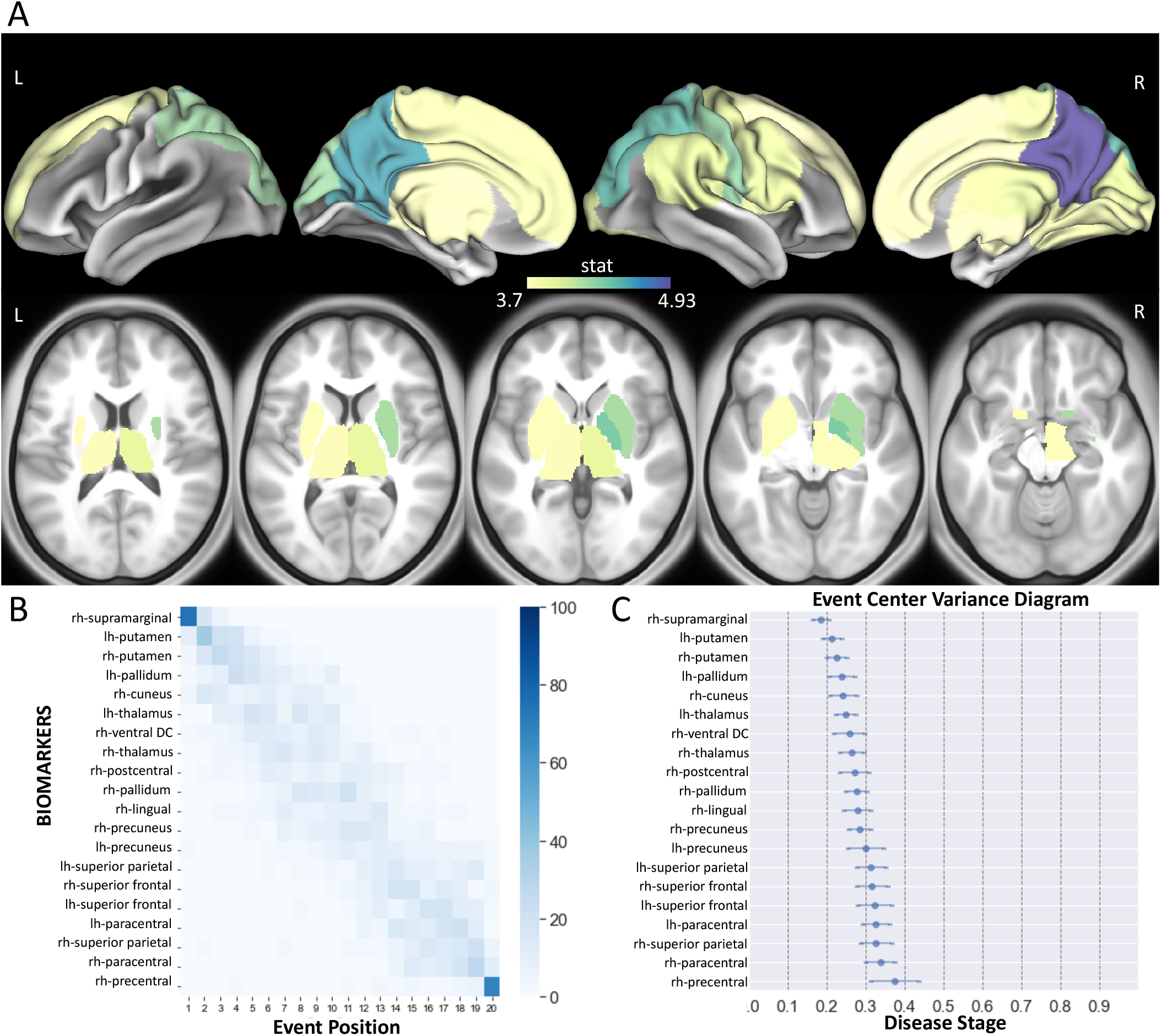
(A) The Student’s t-statistic comparing the log of the T2 FLAIR lesion regional structural disconnectivity between disability groups, displaying the 20 regions having the largest t-statistics (positive t-statistic = structural disconnectivity is greater in individuals with moderate/severe disability). All subjects (n=482) were used for this analysis. (B) Positional variance diagram for disability progression for the same 20 regions. The maximum-likelihood sequence of abnormality is shown on the y-axis (top to bottom). Color intensity in each row indicates positional variance: the darker the color, the higher the confidence of the event position across 100 bootstraps. (C) The event-center variance diagram shows the standard error of estimated abnormality centers. rh/R= right hemisphere and lh/L=left hemisphere.

### PRL and non-PRL Maps and Their Subsequent Disruptions to Structural Connections

As mentioned above, only 363 out of 482 patients had QSM imaging and thus had separate lesion masks for PRL and non-PRL. Figure 3 shows the heat maps of lesion masks of non-PRL (n=363) and PRL (n=156 subjects with ≥ 1 PRL) and their resulting structural disconnectivity maps. Heat maps of the lesion masks show that PRL tended to cluster in periventricular WM (Figure 3, A), compared to non-PRLs that are more widespread throughout the WM (Figure 3, B). The structural disconnectivity due to non-PRL was greater compared to those due to PRL. Note the scale differences in the two modalities are primarily due to the relatively fewer PRL subjects as compared to non-PRL.

**Figure 3:**
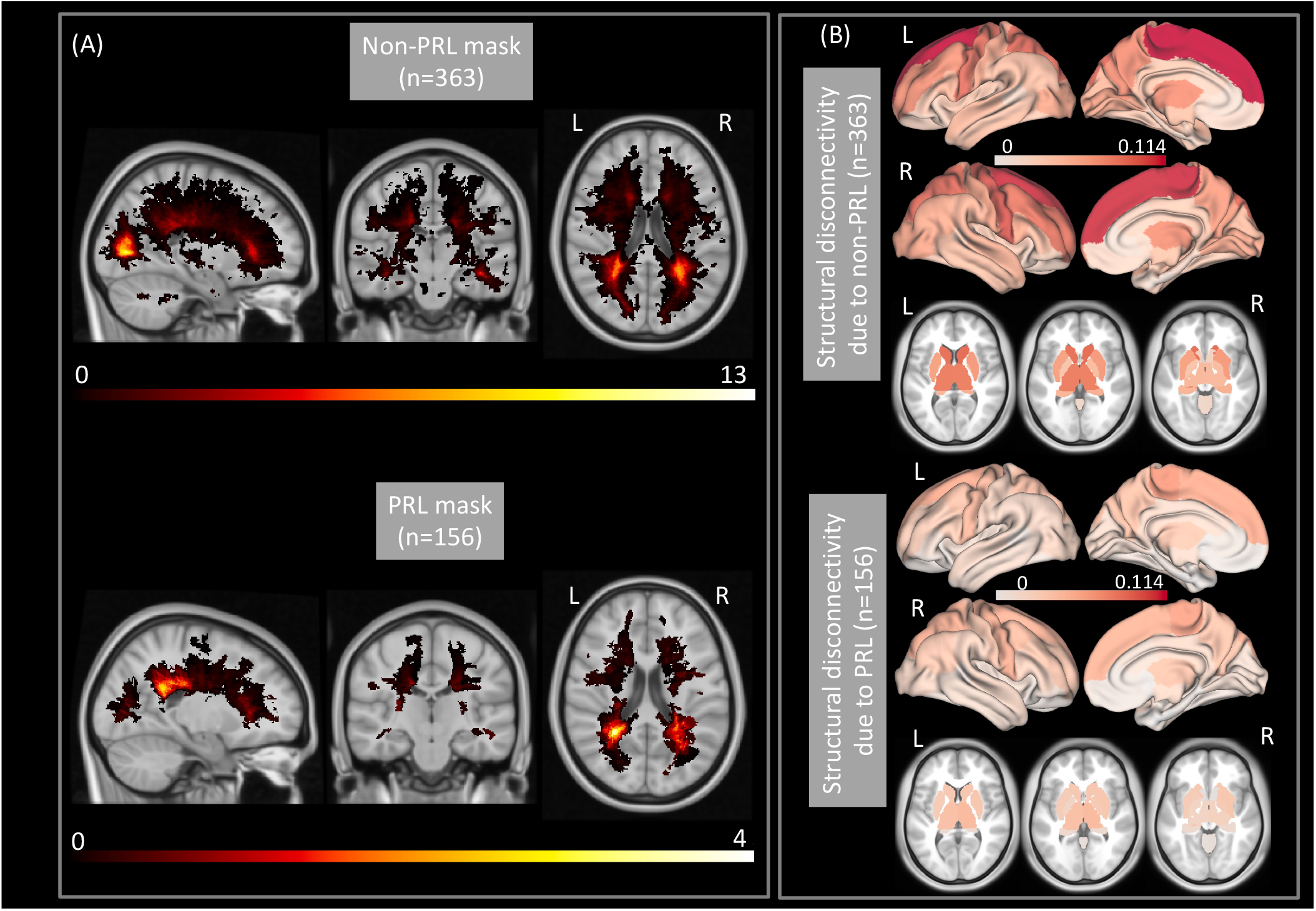
(A) A heatmap of the lesion masks for the non-PRL (T2 FLAIR lesions minus PRL) in 363 subjects and the PRL in 156/363 subjects with ≥ 1 PRL. (B) Regional structural disconnectivity due to the non-PRL and PRL, averaged over their respective cohorts. Darker colors indicate higher structural disconnectivity in the region.

### Sequence of Structural Disconnection due to PRL and non-PRL Across the Spectrum of Disability

We compared the regional structural disconnectivity due to non-PRL or PRL (86 regions x 2 lesion mask types = 172 structural disconnectivity metrics in total) between disability groups in subjects who had QSM imaging and thus had separate lesion masks for PRL and non-PRL (n=363). The 20 regions where the structural disconnectivity was most significantly different (out of these 172 metrics) are presented in Figure 4A, B. The structural disconnectivity metrics were greater in the moderate/severe disability group compared to no/mild disability group across all 20 regions. Only 3/20 of the top metrics were structural disconnections due to PRL. The structural disconnections which were the most significantly different between disability groups were mostly located in the frontal, motor-related, and subcortical regions (Figure 4 A, B). The structural disconnectivity due to PRL and non-PRL in the precentral, postcentral, and paracentral regions were commonly found as greater in the moderate/severe disability group compared to the no/mild disability group. Further, the dEBM results showed that structural disconnectivity due to PRL in the bilateral paracentral gyrus and left precentral gyrus was found to occur first, followed by the structural disconnectivity due to non-PRL in the bilateral caudate, right postcentral gyrus, and right caudal middle frontal gyrus.

**Figure 4:**
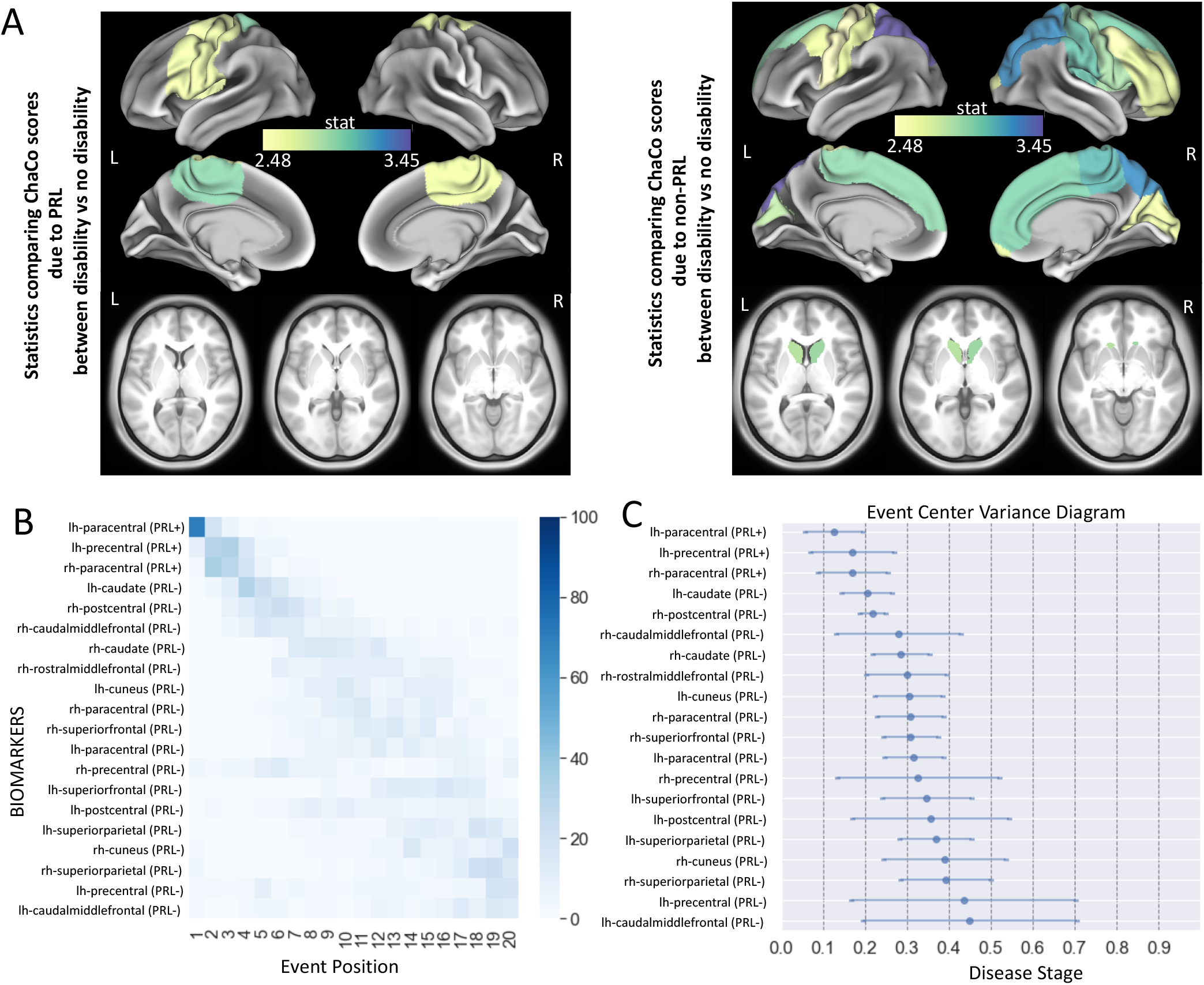
(A) The Student’s t-statistic comparing the log of the regional structural disconnectivity between disability groups; only the 20 regions having the largest t-statistics across both non-PRL and PRL masks are shown. The subjects who had QSM imaging and thus had separate lesion masks for PRL and non-PRL (n=363) were used for this analysis. Darker colors indicate greater t-statistics comparing the structural disconnectivity in the moderate to severe disability group compared to the no to mild disability group. (B) Positional variance diagram across disability progression for the same 20 regions presented in A. The maximum-likelihood sequence of abnormality is shown on the y-axis (top to bottom). Color intensity in each row indicates positional variance: the darker the color, the higher the confidence of the event position across 100 bootstraps. (C) The event-center variance diagram shows the standard error of estimated abnormality centers. rh/R= right hemisphere and lh/L=left hemisphere.

### Sequence of Structural Disconnection due to MS Lesions Across the Spectrum of Cognitive Impairment

Figure 5 shows the t-statistics of the regional (T2 FLAIR lesion-based) structural disconnectivity for the 20 most significantly different regions between CI and CP, as well as the sequence of the regional structural disconnectivity metrics across cognitive impairment. The subjects who had cognitive scores (n=171) were used for this analysis. In all 20 regions, the structural disconnectivity was significantly higher in the CI group compared to the CP group (corrected p-value<0.05), and was most different in the right subcortical and frontal regions. The dEBM results show that structural disconnectivity in right hemisphere subcortical regions, particularly in the right ventral DC, thalamus, and caudate, occurs earlier across cognitive impairment severity.

**Figure 5:**
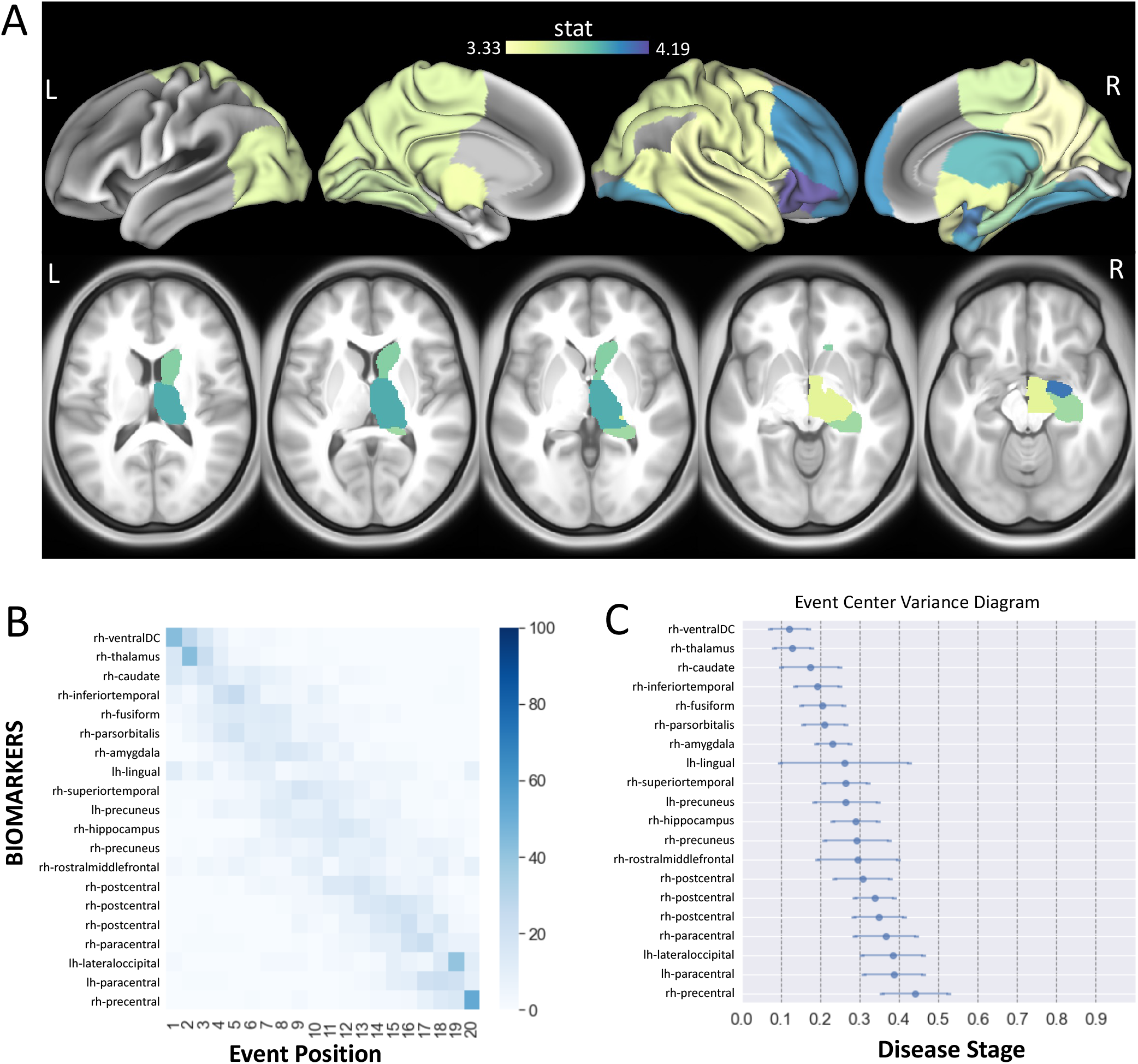
(A) The Student’s t-statistic comparing the log of the T2 FLAIR lesion regional structural disconnectivity between CP and CI groups. The subjects who had cognitive scores (n=171) were used for this analysis. Darker colors indicate greater t-statistics comparing the structural disconnectivity in the CI group compared to the CP group. (B) Positional variance diagram across cognitive impairment severity for the same 20 regions presented in A. The maximum-likelihood sequence of abnormality is shown on the y-axis (top to bottom). Color intensity in each row indicates positional variance: the darker the color, the higher the confidence of the event position across 100 bootstraps. (C) The event-center variance diagram shows the standard error of estimated abnormality centers. rh/R= right hemisphere and lh/L=left hemisphere.

As mentioned in Methods section, patients were assigned the CI group when at least 2 out of 3 cognitive tests were identified as below normal. As a post-hoc analysis, we reduced this requirement to only 1 out of 3 cognitive tests being abnormal for assignment to the CI group. We found similar results wherein the structural disconnectivity in subcortical and frontal, along with visual regions occur in the earlier stages across cognitive impairment severity.

### Sequence of Structural Disconnection due to PRL and non-PRL Across the Spectrum of Cognitive Impairment

We also investigated the sequence of PRL and non-PRL-based structural disconnectivity across cognitive impairment severity in all patients with cognitive scores (n=171). Our results showed that structural disconnectivity due to non-PRL occurs at earlier stages compared to structural disconnectivity due to PRL across cognitive impairment severity. Similar to our results with T2 FLAIR lesions, non-PRL structural disconnectivity in subcortical regions, particularly in the right ventral DC, caudate, and thalamus, occurs early in the event sequence.

## DISCUSSION

In this study, we used dEBM applied to cross-sectional data in pwMS to investigate the sequence of regional structural disconnectivity across the spectrum of disability and cognitive impairment. We computed structural disconnectivity metrics using the NeMo tool and overall T2 FLAIR lesions as well as these lesions separated into QSM-hyperintense paramagnetic rim (PRL) or no rim (non-PRL). We showed that structural disconnectivity due to T2 FLAIR lesions in the ventral attention and subcortical networks, in particular within the supramarginal and putamen, occurs earlier in the sequence of disability (i.e., mild to severe). We also showed that PRL-based structural disconnectivity in motor-related regions occurs earlier in the disability event sequence, followed by non-PRL-based structural disconnectivity in the caudate and postcentral gyrus. T2 FLAIR lesion-based structural disconnectivity in subcortical regions, including the thalamus, occurs earliest in the sequence of cognitive impairment (i.e. preserved to impaired).

### Structural disconnectivity in subcortical regions is central to disability and cognitive impairment

The distribution of MS lesions is not uniform across the brain. Histopathological studies have shown that MS lesions tend to more frequently appear in periventricular, juxtacortical, and subpial locations^1^; however, lesion location and size are quite heterogeneous which contributes to the widely varying physical and cognitive impairment among pwMS ^31–34^.

Subcortical regions have been shown to play a large role in MS, as atrophy in this region has been associated with motor and cognitive impairment as well as fatigue^35,36^. In alignment with these findings, one of our recent studies demonstrated that increased structural disconnectivity due to PRL and non-PRL in the thalamus was associated with greater disability in pwMS^5^. Moreover, other studies have shown that atrophy in subcortical regions, particularly in the thalamus, pallidum, putamen, and cuneus occurs as an early event in relation to disability progression in MS.^2,3^ These results are consistent with the findings of our study that uses structural disconnectivity metrics which were previously shown to be associated with atrophy in certain subcortical regions.^37^

Only one study investigated the sequence of regional/network-based structural and functional changes across cognitive impairment severity in MS^3^. This study used a different cognitive test than our study (i.e. Brief Repeatable Battery of Neuropsychological Tests) to assign subjects into CI, mildly CI, and CP groups and investigated the sequence of the events across the spectrum of cognitive impairment. Thalamic atrophy was found to occur second, just after increased functional centrality in the default mode network, as cognitive impairment became more severe. This is in concordance with our results that show structural disconnection in the right hypothalamus (ventral DC) and right thalamus occurs earliest across the spectrum of cognitive impairment. Thalamic lesions are very common in MS; 71% of patients have thalamic lesions on 7T MRI^38^ and larger thalamic lesions have been associated with greater cognitive impairment^39^. Altogether, these results support the commonly reported finding that damage to the thalamus, either in gray matter or connected WM, plays an important role in both physical and cognitive impairment in MS.

### Structural disconnectivity due to PRL occurs earlier in the spectrum of disability severity

While the impact of lesion location on disability type/severity has been previously studied in MS ^5,31,40^, there are fewer studies that attend to the role of chronic lesion subtype, i.e. chronic active (PRL) vs inactive (non-PRL). Our previous study found that structural disconnectivity from PRL more accurately predicted disability compared to structural disconnectivity from non-PRL^5^. Moreover, increased PRL-based structural disconnectivity in the thalamus and cerebellum was commonly found to be associated with worse disability. However, the sequence of PRL vs. non-PRL’ resulting structural disconnections across disability severity had not been studied. Our study shows that structural disconnectivity due to PRL, particularly in motor-related regions such as bilateral paracentral and left precentral, occurs earlier across the spectrum of disability severity. It may be for this reason that chronic active lesions, although the far minority of lesions, demonstrate an association with clinical disability^41,42^, and these results suggest that they may be an early biomarker of disease progression. It has been previously shown that the neurodegeneration due to MS lesions may impact both the axonal terminal and the cell soma, which may then lead to atrophy in the connected GM regions.^43^ WM lesion volume was shown to be associated with GM atrophy at both global and regional level.^44–46^ Thus, the structural disconnectivity due to PRL in the motor-related regions might cause atrophy in these regions. Our findings align with previous evidence that physical dysfunction and atrophy in the sensorimotor cortex are related ^47–50^. Interestingly, PRL did not have the same relevance when evaluating cognitive impairment as compared to EDSS and we suspect this difference may relate to relatively small number of CI patients in combination with a relatively rare occurrence of a PRL. PRLs have been associated with cognitive impairment ^41,42^, thus this should be repeated in a larger sample of patients with cognitive testing.

### Discriminative event-based modeling in pwMS

DEBM allows investigation of the sequence of biomarker abnormality across disability/impairment progression using large cross-sectional neuroimaging datasets, which are easier to collect than longitudinal data. This approach has been applied to data from pwMS to investigate the sequence of regional atrophy ^2^ as well as structural, functional, and cognitive changes^3^. The latter study used structural connectivity measured with diffusion MRI^3^, which stands in contrast to our approach that estimates the structural disconnection from the lesion masks and the NeMo Tool. In addition to its wider clinical applicability, our approach to measure structural disconnectivity has also been shown to be a better predictor of disability compared to structural and functional connectivity metrics computed with advanced imaging techniques^8^. Lastly, our study did not use a-priori selected regions but rather included all cortical, subcortical, and cerebellar regions before statistical selection of the regions to include in the dEBM. This data-driven approach is advantageous as it is informed by the computationally intensive use of the existing dataset, therefore provides less biased conclusions from the data compared to hypothesis-driven approach.

### Limitations

A limitation of our study is that the regional structural disconnectivity metrics were computed by considering only the lesion size and location. As various pathologies such as myelin and/or axonal content cannot be discerned on T2 FLAIR, the differential impact of specific lesions’ pathologies on structural connections was not considered. In addition, the regional sequence of the structural disconnections was not investigated for different phenotypes of MS such as CIS, RRMS, SPMS, and PPMS. A future study could investigate if the sequence of the structural disconnections differs for different phenotypes might be beneficial, however traditional clinical classifiers may not be relevant^51^. Presence or absence of treatment and type of treatments likely contribute to disability progression and the appearance of new lesions; this information was not included in the dEBM model. A study that includes a similar amount of patients in each treatment category should be performed. Finally, the control subjects in the NeMo Tool ranged mostly from 21-35, which is younger than many of the pwMS in our study; any aging effects on WM or structural connectivity in the pwMS would not be considered in the structural disconnection metrics.

## Conclusion

This is the first study to investigate the sequence of lesion-based regional structural disconnectivity across the spectrum of disability and cognitive impairment severities in pwMS. We found that structural disconnections in subcortical regions, including the thalamus, occur earlier in the spectrum of disability and cognitive impairment. We also found that PRL-based structural disconnectivity in motor-related regions occurs earliest in disability progression. This study provides a better understanding of how structural disconnectivity due to T2 FLAIR lesions and PRL/non-PRL deferentially impacts disability and cognitive impairment in MS. A deeper understanding of the regional pattern of structural disconnection due to different types of lesions may help in identifying the early biomarkers of disability, creating more accurate prognoses and perhaps personalize treatment selection to reduce the burden of MS.

## Abbreviations

BVMT: Brief Visuospatial Memory Test
BH: Benjamini-Hochberg
CI: cognitively impaired
CP: cognitively preserved
CIS: clinically isolated syndrome
CVLT: California Verbal Learning Test
dEBM: discriminative event-based modeling
dMRI: diffusion MRI
EDSS: expanded disability status scale
GM: gray matter
LST: lesion segmentation tool
MS: multiple sclerosis
NeMo: network modification
PPMS: primary progressive MS
PRL: paramagnetic rim lesions
pwMS: people with multiple sclerosis
SPMS: secondary progressive
SDMT: Symbol Digit Modalities Test
QSM: quantitative susceptibility mapping
WM: white matter

## ACKNOWLEDGEMENTS

C.T. helped for post-processing, designed the study, performed the statistical analysis, and wrote the manuscript. E.O. helped with post-processing and reviewed the article. K.J. performed pre- and post-processing of MRI data and reviewed the article. E.D. helped for data collection and creating lesion masks. U.K. manually edited the lesion masks, collected patient data, and edited the article. M.M. helped for data collection and creating lesion masks. N.Z. helped for data collection and creating lesion masks. N.M. helped with creating lesion masks, post-processing, and reviewing the article. N.S. helped for data collection and creating lesion masks. T.N. helped for image data processing and analysis. S.G. designed and supervised the study, collected the data, and edited the article. A.K. designed and supervised the study, collected the data, and edited the article.

## Data/code availability statement

The deidentified data that support the findings of this study are available upon reasonable request from the corresponding author. The codes which were used to perform the dEBM analysis are publicly available (https://github.com/EuroPOND/pyebm).

## Ethics statement

All studies were approved by an ethical standards committee on human experimentation, and written informed consent was obtained from all patients.

